# Systematic misperceptions of 3D motion explained by Bayesian inference

**DOI:** 10.1101/149104

**Authors:** Bas Rokers, Jacqueline M. Fulvio, Jonathan Pillow, Emily A. Cooper

## Abstract

People make surprising but reliable perceptual errors. Here, we provide a unified explanation for errors in the perception of three-dimensional (3D) motion. To do so, we characterized the retinal motion signals produced by objects moving with arbitrary trajectories through arbitrary locations in 3D. Next, we developed a Bayesian model, treating 3D motion perception as optimal inference given sensory noise and the geometry of 3D viewing. The model predicts a wide array of systematic perceptual errors, that depend on stimulus distance, contrast, and eccentricity. We then used a virtual reality (VR) headset as well as a standard 3D display to test these predictions in both traditional psychophysical and more naturalistic settings. We found evidence that people make many of the predicted errors, including a lateral bias in the perception of motion trajectories, a dependency of this bias on stimulus contrast, viewing distance, and eccentricity, and a surprising tendency to misreport approaching motion as receding and vice versa. In sum, we developed a quantitative model that provides a parsimonious account for a range of systematic misperceptions of motion in naturalistic environments.

## Introduction

The accurate perception of visual motion is critical for everyday behavior. In the natural environment, motion perception involves determining the 3D direction and speed of moving objects, based on both retinal and extra-retinal sensory cues. In the laboratory, a large number of studies have reported systematic biases in the perception of 3D motion, despite the availability of many such cues (Fulvio, Rosen, & Rokers, 2015; Harris & Dean, 2003; Harris & Drga, 2005; Lages, 2006; Rushton & Duke, 2007; Welchman, Lam, & Bülthoff, 2008; Welchman, Tuck, & Harris, 2004). These perceptual errors may contribute to behavioral failures in real world scenarios, such as catching projectiles (Peper, Bootsma, Mestre, & Bakker, 1994) and driving under foggy conditions (Pretto, Bresciani, Rainer, & Bülthoff, 2012; Shrivastava, Hayhoe, Pelz, & Mruczek, 2010; Snowden, Stimpson, & Ruddle, 1998). Here we ask if a range of systematic errors in 3D motion perception can be understood as the consequence of 3D viewing geometry and reasonable prior expectations about the world.

Bayesian observer models are a strong candidate for addressing this question. They provide a straightforward rule for the optimal combination of incoming sensory evidence with prior knowledge. The Bayesian framework has successfully explained a variety of perceptual phenomena (Girshick, Landy, & Simoncelli, 2011; Knill, 2007; Knill & Richards, 1996), including systematic biases in 2D motion perception (Weiss, Simoncelli, & Adelson, 2002). Specifically, when visual input is unreliable (for example, when a stimulus has low contrast), observers systematically underestimate the speed of visual motion in the fronto-parallel plane: low-contrast patterns appear to move more slowly than otherwise equivalent high contrast patterns (Stone & Thompson, 1992; Thompson, 1982). This misperception, along with several other seemingly unrelated phenomena in motion perception, can be elegantly accounted for by a Bayesian model that incorporates a prior assumption that objects in the world tend to move slowly (Stocker & Simoncelli, 2006; Weiss et al., 2002).

Errors occur in the domain of 3D motion perception as well. For example, observers systematically overestimate angle of approach in 3D, such that objects moving towards the head are perceived as moving along a path that is more lateral than the true trajectory (we call this a ‘lateral bias’) (Harris & Dean, 2003; Harris & Drga, 2005; Lages, 2006; Rushton & Duke, 2007; Welchman et al., 2008, 2004). Bayesian models of 3D motion perception, also assuming a slow motion prior, have been shown to account for this bias (Lages, 2006; Welchman et al., 2008). However, existing models are restricted to specific viewing situations (stimuli in the midsagittal plane), and have been tested using tasks and stimuli that limit the kind of perceptual errors that can be observed.

Here, we provide a model of 3D motion perception that can predict systematic errors in the perception of motion, extending to arbitrary stimulus locations and naturalistic tasks. First, we derived a general Bayesian model for 3D motion estimation from retinal motion cues that does not depend on any specific viewing situation. The model generates predictions of perceived motion for stimuli in arbitrary locations in 3D space. The full model reveals that — contrary to previous conclusions — motion-in-depth estimation is not fundamentally less reliable than lateral motion estimation (Tyler, 1971; Welchman et al., 2008). Instead, the relative increase in sensory uncertainty for motion-in-depth derives from the geometry of typical viewing situations in the laboratory.

Second, we used this model to quantify the relationship between stimulus contrast, stimulus location, and perceptual errors. Like previous models, the current model predicts a lateral bias in perception of motion trajectories towards the head, but also a clear effect of viewing distance. Since many different visual cues to 3D motion are present under natural viewing conditions, and the predicted errors are derived from only one visual cue (binocular retinal motion), we were particularly interested in determining whether these perceptual errors occur under conditions where a rich array of sensory cues is available. We therefore conducted experiments in a virtual reality (VR) environment, which provided an intuitive response paradigm and a rich array of cues to 3D motion. We established that the lateral bias does occur in a three-dimensional environment that simulates these naturalistic conditions, and that the bias is modulated by distance and contrast as predicted by the model.

Third, we examined a recently identified perceptual phenomenon in which the direction of motion-in-depth (but not lateral motion) is fundamentally misreported: approaching motion is reported to be receding and vice versa (Fulvio, Rosen, & Rokers, 2015). We established that the model also predicts these motion-in-depth misreports, and identified the relationship between these misreports and viewing conditions. We subsequently confirmed the presence of these misreports in the VR paradigm described above, and in a separate experiment using a conventional stereoscopic display.

Finally, we generated model predictions specifically for perceived motion trajectories originating ‘off to the side’, outside of the observer’s midsagittal plane. We subsequently presented motion stimuli with the use of the wide-angle VR display, and confirmed the predicted lawful relationship between perceived motion trajectory and stimulus eccentricity. We thus provide a unified account of multiple perceptual phenomena in 3D motion perception, showing that geometric considerations, combined with optimal inference under sensory uncertainty, explain these systematic and, at times, dramatic misperceptions.

## Methods

Here, we describe the methods for three experimental studies used to test a range of perceptual predictions made by the Bayesian model.

### Experiment 1

The goal of Experiment 1 was to test the model predictions regarding the effects of viewing distance and stimulus contrast on perceptual errors (lateral bias and direction misreports) in the midsagittal plane using a naturalistic VR paradigm.

#### Participants

Sixty-eight college-aged members of the University of Wisconsin-Madison community (38 female, 30 male) gave informed consent to complete the study, and 47 (26 female, 21 male) successfully completed all parts of the experiment. The participants that did not complete the study either had difficulty understanding the task, perceiving depth in the display, or wearing glasses inside the VR head-mounted display system. The experiment was carried out in accordance with the guidelines of The University of Wisconsin–Madison Institutional Review Board. Course credits were awarded in exchange for participation.

All participants had normal or corrected-to-normal vision and were screened for intact stereovision using the RANDOT Stereotest (Stereo Optical Company, Inc., 2011). To qualify for the study, participants were required to accurately identify all of the shapes in the RANDOT Form test, to identify the location of at least 5 out of 10 targets in the RANDOT Circle test, and to pass the suppression check. Although all participants passed the tests at these criteria, those with lower scores on the Form test (i.e., with a score of 5 or 6) were more likely to terminate their participation early (~50% of those who consented but did not complete the study).

#### Apparatus

The experiment was controlled by Matlab and the Psychophysics Toolbox (Brainard, 1997; Kleiner, Brainard, Pelli, Ingling, & Murray, 2007; Pelli, 1997) on a Macintosh computer and projected through the Oculus Rift Development Kit 1 (DK1) (www.oculusvr.com), which was calibrated using standard gamma calibration procedures. The Oculus Rift DK1 is a stereoscopic head-mounted VR system with an 18 cm LCD screen embedded in the headset providing an effective resolution of 640x800 pixels per eye with a refresh rate of 60 Hz. The horizontal field of view is over 90 deg (110 deg diagonal). Also embedded within the headset is a 1000 Hz Adjacent Reality Tracker that relies upon a combination of gyros, accelerometers, and magnetometers to measure head rotation along the yaw, pitch, and roll axes with a latency of 2 ms. Note that translations of the head are not tracked by the device. Participants used a wireless keyboard to initiate trials and make responses.

#### Stimulus & Procedure

In a series of trials, participants were asked to indicate the perceived direction of motion of a target sphere that moved with a constant velocity in the virtual environment. The stimuli were presented in the center of a virtual room (3.46 m in height, 3.46 m in width, and 14.4 m in depth). The virtual wall, ceiling, and floor were all mapped with different textures. These textures were included to facilitate better judgment of distances throughout the virtual space and the relative positions of the stimuli (**Figure 1a**).

**Figure 1.**
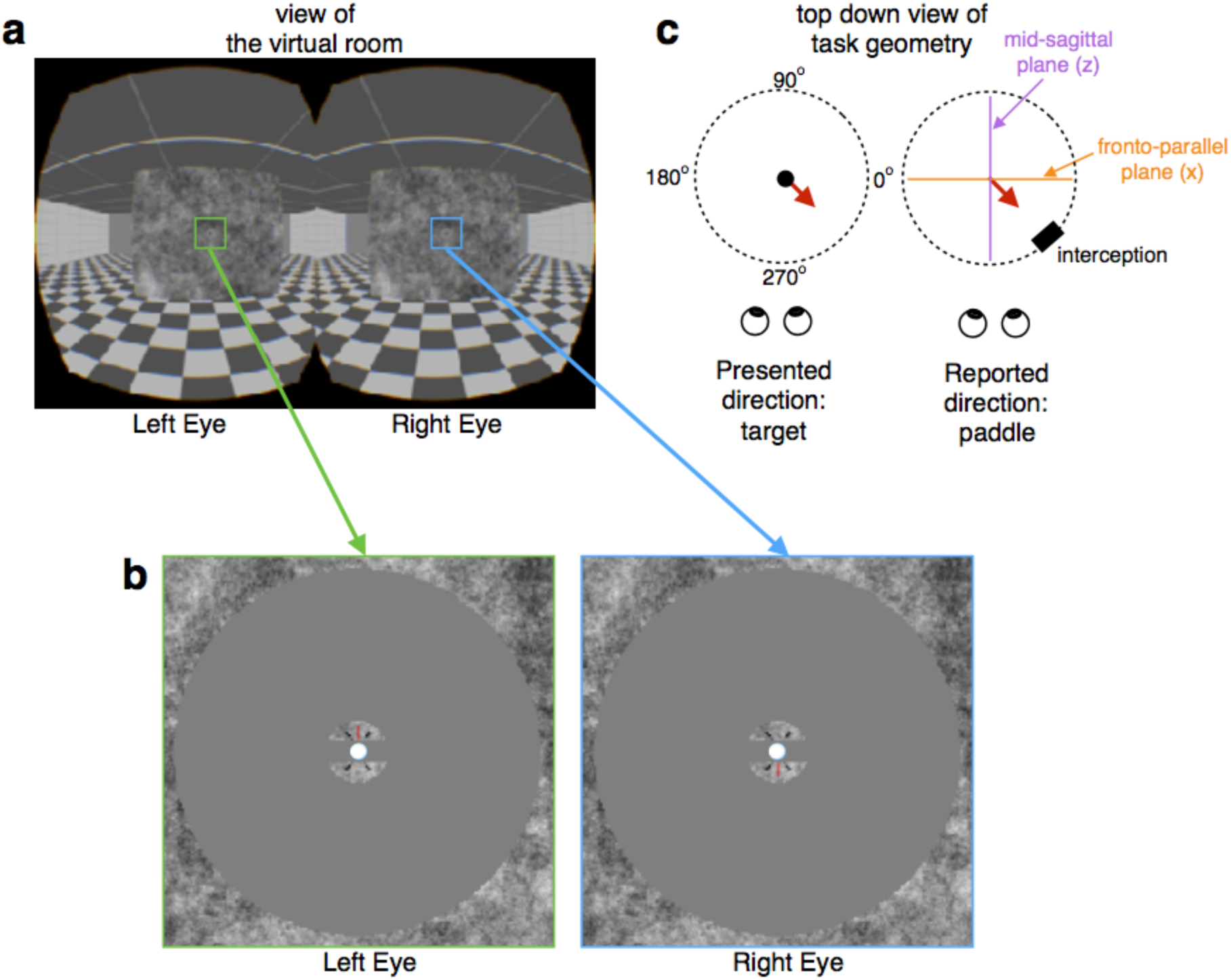
Stimulus and procedure for Experiment 1. **(a)** Participants wore a head-mounted display and viewed a stereoscopic virtual room with a planar surface in the middle. **(b)** Zoomed in views of the left and right eye’s images show the critical aspects of the stimulus. Participants fixated nonius lines in the center of a circular aperture, and a virtual target (white sphere) appeared inside the nonius lines. **(c)** The target moved at a constant velocity in a random direction within the xz-plane (Presented direction). Afterwards, participants positioned a virtual paddle such that it would intersect the trajectory of the target (Reported direction). The setting denoted by the black paddle in this example would result in a successful target interception.

The stimuli were otherwise similar to those used in Fulvio et al. (2015). In the center of the virtual room, there was a planar surface with a circular aperture (7.5° in radius). The planar surface was mapped with a 1/f noise pattern that was identical in both eyes to aid vergence. In addition, nonius lines were embedded within a small 1/f noise patch near the center of the aperture. All stimulus elements were anti-aliased to achieve subpixel resolution. The background seen through the aperture was mid-gray (**Figure 1b**).

The planar surface was positioned in the room at one of two viewing distances from the observer’s location: 90 cm (n=15 participants) and 45 cm (n=32 participants). Participants were instructed to fixate the center of the aperture. However, participants were free to make head movements, and when doing so, the display updated according to the viewpoint specified by the yaw, pitch, and roll of the participant’s head. Translations of the head did not impact the display, such that stimulus viewing distance remained constant.

On each trial, a white sphere (‘target”) of size 0.25° in diameter appeared at the center of the aperture and then followed a trajectory defined by independently chosen random speeds in the x (lateral) direction and the z (motion-in-depth) direction, with no change in y (vertical direction) before disappearing. The motion trajectory always lasted for 1 s. Velocities in x and z were independently chosen from a 2D Gaussian distribution (M = 0 cm/s, SD = 2 cm/s) with imposed cut offs at 6.1 cm/s and -6.1 cm/s. The independently chosen speeds resulted in motion trajectories whose directions spanned the full 360° space (**Figure 1c,** left side). Thus, the target came toward the participant (‘approaching’), and moved back behind fixation away from the participant (‘receding’) on approximately 50% of trials each. It is important to note that since x- and z- motion were chosen randomly and independently, the amount of perceived lateral movement on each trial did not carry information about the amount of motion-in-depth and vice versa. The target was rendered under perspective projection, so that both monocular (looming) and binocular cues to motion-in-depth were present.

Participants indicated the perceived target trajectory using a “3D pong” response paradigm (Fulvio et al., 2015). After the target disappeared, a 3D rectangular block (‘paddle’), whose faces also consisted of a 1/f noise pattern, appeared at the edge of the aperture. The paddle dimensions were 0.25 cm x 0.5 cm x 0.25 cm. Participants were asked to extrapolate the target’s trajectory and adjust the paddle’s position such that the paddle would have intercepted the target if the target had continued along its trajectory. The paddle’s position could be adjusted along a circular path that orbited the fixation point in the x-z plane using the left and right arrow keys of the keyboard (**Figure 1c,** right side). As the participant moved the paddle through the visual scene, it was rendered according to the rules of perspective projection. Thus, the stimuli were presented and the responses were made in the same 3D space. By asking participants to extrapolate the trajectory, we prevented participants from setting the paddle to a screen location that simply covered the last seen target location. We did not ask participants to retain fixation during the paddle adjustment phase of the trial. When the participant was satisfied with the paddle setting, they resumed fixation and pressed the spacebar to initiate a new trial. A stereoscopic movie demonstrating the general procedure is included in the Supplementary Material (**Movie S1**).

The target was presented at one of three Michelson contrast levels: 100% (high), 15% (mid), and 7.5% (low), which were counterbalanced and presented in pseudorandom order.

Participants carried out 10-15 practice trials in the presence of the experimenter to become familiar with the task. All participants completed the experimental trials in one session. No feedback was provided for either the practice or experimental trials. Participants completed 225 trials on average.

#### Data Analysis

To examine biases in the perceived direction of motion, we computed the mean angular error for each participant for each unique stimulus condition (viewing distance and target contrast). Errors were calculated as the angular distance of the reported direction relative to the presented direction in the xz-plane. We analyzed the data to determine whether this angular error tended to be towards the fronto-parallel plane (lateral bias) or towards the mid-sagittal plane (medial bias) (See **Figure 1c)**. We assigned positive values to medial errors and negative values to lateral errors such that the average would indicate the overall directional bias.

To examine the frequency of motion direction errors, we computed the percentage of trials on which paddle settings were made on the opposite side of either the fronto-parallel plane or the mid-sagittal plane. Responses on the opposite side of the fronto-parallel plane (approaching vs. receding) were considered depth direction confusions. Responses made on the opposite side of the mid-sagittal plane (leftward vs. rightward) were considered lateral direction confusions.

Statistical effects were tested through an analysis of variance (ANOVA) evaluated on generalized linear model fits. The model incorporated target contrast as a fixed effect (all subjects experienced all three levels) and viewing distance as a random effect (different groups of subjects performed the task for one of the two viewing distances). The model intercepts were included as random subject effects. Follow-up tests consisted of Bonferroni-corrected t-tests for multiple comparisons.

### Experiment 2

To examine whether the motion direction confusions measured in Experiment 1 were particular to the VR set up, we compared these results to a second experiment conducted on a traditional stereoscopic display.

#### Participants

Three adults participated in the experiment. All had normal or corrected-to-normal vision. One participant (male, age 23) was naive to the purpose of the experiment and had limited psychophysical experience. The remaining two participants (the authors JP and BR, males aged 34–35) had extensive psychophysical experience. The experiment was undertaken with the written consent of each observer, and all procedures were approved by The University of Texas at Austin Institutional Review Board.

#### Apparatus

The experiment was performed using a similar setup to Experiment 1, however in this case the stimuli were presented on two 35.0 x 26.3 cm CRT displays (ViewSonic G90fB, one for each eye; 75 Hz, 1280 x 1024 pixels) at a single viewing distance of 90 cm (21.2 x 16.3 deg of visual angle). Left- and right-eye half-images were combined using a mirror stereoscope. The luminance of the two displays was linearized using standard gamma-correction procedures, and the mean luminance was 50.6 cd/m^2^.

#### Stimulus & Procedure

As in Experiment 1, all stimuli were presented within a circular mid-gray aperture (1 deg radius) that was surrounded by a 1/f noise texture at the depth of the fixation plane (90 cm) to help maintain vergence. No virtual room was present. Additionally, a small square fixation point was placed at the center of the display. The fixation point was surrounded by horizontal and vertical nonius lines, and was placed on a circular 0.1 deg radius 1/f noise pattern.

Rather than a single target, a field of randomly positioned dots moving in the xz-plane was presented on each trial. The positions of the dots were constrained to a single plane fronto-parallel to the display (i.e., perpendicular to the observer’s viewing direction). The initial disparity of the plane was consistent with a distance of 93 cm (3 cm behind the fixation plane). The plane then moved for 500 ms with an x and z velocity independently and uniformly sampled from a -4 to 4 cm/s interval, corresponding to a maximum possible binocular disparity of 0.21 deg (uncrossed) relative to the fixation plane.

Each moving dot had a 200 ms limited lifetime to prevent tracking of individual stimulus elements. Dot radius was 0.11 cm and dot density was ~74 dots/deg^2^. Both dot size and dot density changed with distance to the observer according to the laws of perspective projection. Dots were presented at one of three Weber contrast levels (7.5, 15, or 60%). Half of the dots were darker, and the other half of the dots were brighter than the mid-gray background.

The stereoscope was initially adjusted so that the vergence demand was appropriate for the viewing distance and given a typical interocular distance. Prior to each session, each participant made further minor adjustments so that the nonius lines at fixation were aligned both horizontally and vertically, and vergence was comfortable. Participants were instructed to maintain fixation for the duration of each experimental trial.

Trials proceeded as described for Experiment 1, except that participant responses were made differently. After the dots disappeared, a circle and a line were presented on screen, where one of the line endpoints was fixed to the center of the circle and the participant could adjust the other line endpoint with a computer mouse. Participants were instructed to treat this as a top-down view of the stimulus (see **Figure 1c**), and to adjust the line such that the angle was consistent with the trajectory of the dots. We verified in pilot experiments that this method produced consistent, reproducible estimates. As in Experiment 1, no feedback concerning performance was provided.

#### Data Analysis

Data were analyzed in the same manner as Experiment 1.

### Experiment 3

To test model predictions for stimuli presented at locations ‘off to the side’ – away from the midsagittal plane – we conducted a third experiment using the same virtual reality display as described in Experiment 1.

#### Participants

Twenty-two college-aged members of the University of Wisconsin-Madison community gave informed consent to complete the study. One participant did not complete the study because of difficulty in perceiving depth in the display, despite passing the stereovision screening (see below). The remaining 21 participants completed all aspects of the experiment. The experiment was carried out in accordance with the guidelines of The University of Wisconsin–Madison Institutional Review Board. Course credits were awarded in exchange for participation. All participants had normal or corrected-to-normal vision and were screened for intact stereovision using the RANDOT Stereotest (Stereo Optical Company, Inc., 2011) in order to meet the criteria outlined in Experiment 1 above.

#### Apparatus

The apparatus was the same as that described in Experiment 1.

#### Stimulus & Procedure

The stimulus and procedure were the similar to Experiment 1, with the exception that the planar surface in the center of the room had three circular apertures rather than just one. As in Experiment 1, one of the apertures appeared at the center of the planar surface directly in front of the participants (7.5 deg radius). The other two apertures were located 20 degrees to the left and right of the central location. These two apertures were slightly larger (10.5 deg radius) in order to ensure adequate visibility of the stimulus. All three apertures appeared on every trial, and the background seen through the aperture was black, which increased the contrast of the stimuli, and further improved visibility in the periphery (**Figure 1b**).

The planar surface was positioned in the virtual room at 45 cm from the participants. Participants were instructed to fixate the center of the central aperture on every trial, even when a target appeared at one of the peripheral locations. This instruction served to minimize head rotation, not eye-movements per se. As we will see model predictions critically depend on stimulus location in 3D space (relative to the observer), not stimulus position on the retina.

On each trial, a white sphere (‘target”) appeared at the center of one of the three apertures randomly and counter-balanced across trials. To ensure that the peripheral targets were clearly visible while participants fixated the central aperture, the peripheral targets were rendered with a diameter of 0.5 deg, versus the 0.25 deg in the central location. The target was presented with full, 100% contrast – i.e., white on a black background. All other aspects of the target’s motion were identical to Experiment 1 above.

Participants indicated the perceived target trajectory using a “3D pong” response paradigm as in Experiment 1 at each of the three aperture locations. They were free to move their eyes to the three apertures during the response phase of the trial. Participants carried out 10-15 practice trials in the presence of the experimenter to become familiar with the task. All participants completed the experimental trials in one session. No feedback was provided for either the practice or experimental trials. All participants completed 360 experimental trials.

#### Data Analysis

Data were analyzed in the same manner as Experiment 1.

## Results

### Geometric explanation of biases in 3D motion perception

The Bayesian brain hypothesis posits that perception is a probabilistic process in which the perception of the physical world is dictated by a probability distribution called the posterior (Knill & Richards, 1996; Knill & Pouget, 2004). This posterior specifies the conditional probability of the physical stimulus (*s*) given a sensory response (*r*) — denoted *P*(*s* | *r*). The posterior is determined by the product of two probabilistic quantities known as the likelihood and the prior. The likelihood is the conditional probability of the observed sensory responses (*r*) given the physical stimulus (*s*), considered as a function of *s*. The likelihood, written *P*(*r* | *s*), characterizes the information that neural responses carry about the sensory stimulus. Increased sensory uncertainty, due to ambiguity or noise in the external world, or internal noise in the sensory system, manifests as an increase in the width of the likelihood. The prior *P*(*s*) represents the assumed probability distribution of the stimulus in the world. The prior may be based on evolutionary or experience-based learning mechanisms. The relationship between posterior, likelihood, and prior is given by Bayes’ rule, which states that:

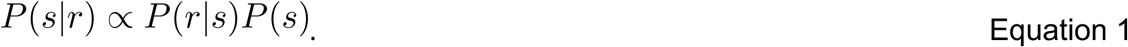

When sensory uncertainty is high, the likelihood is broad and the prior exerts a relatively large influence on the posterior, resulting in percepts that are systematically more biased towards the prior (but see Wei & Stocker, 2015). That is, the visual system relies on prior assumptions when current information is unreliable. Misperceptions will inevitably occur when actual stimulus properties diverge from these prior assumptions, particular when sensory uncertainty is high.

Here we apply this Bayesian framework to the problem of 3D motion perception. While the derivation of the posterior distribution for 3D motion is lengthy, we first provide an intuition by examining a simple diagram illustrating the consequences of perspective projection on retinal signals to 3D motion. First, we consider that light from moving objects in the world will project through the optics of the eye and cast a pattern with a particular angular velocity on the retina. This is illustrated in **Figure 2a**. A simplified top-down diagram illustrates the left and right eyes of an observer (projections are shown for the left eye only; the right eye is for reference). Two linear motion vectors are illustrated in orange and green. The vectors have the same length, indicating the same speed in the world, but they move in different directions, either in depth (towards the observer, shown in green) or laterally (to the left, shown in orange). Of course, the angular velocity signal in either eye alone does not specify the direction of motion in the world. While these signals do constrain the possible trajectories, estimates of 3D motion critically depend on the *relationship* of the retinal velocity signals between the two eyes.

**Figure 2.**
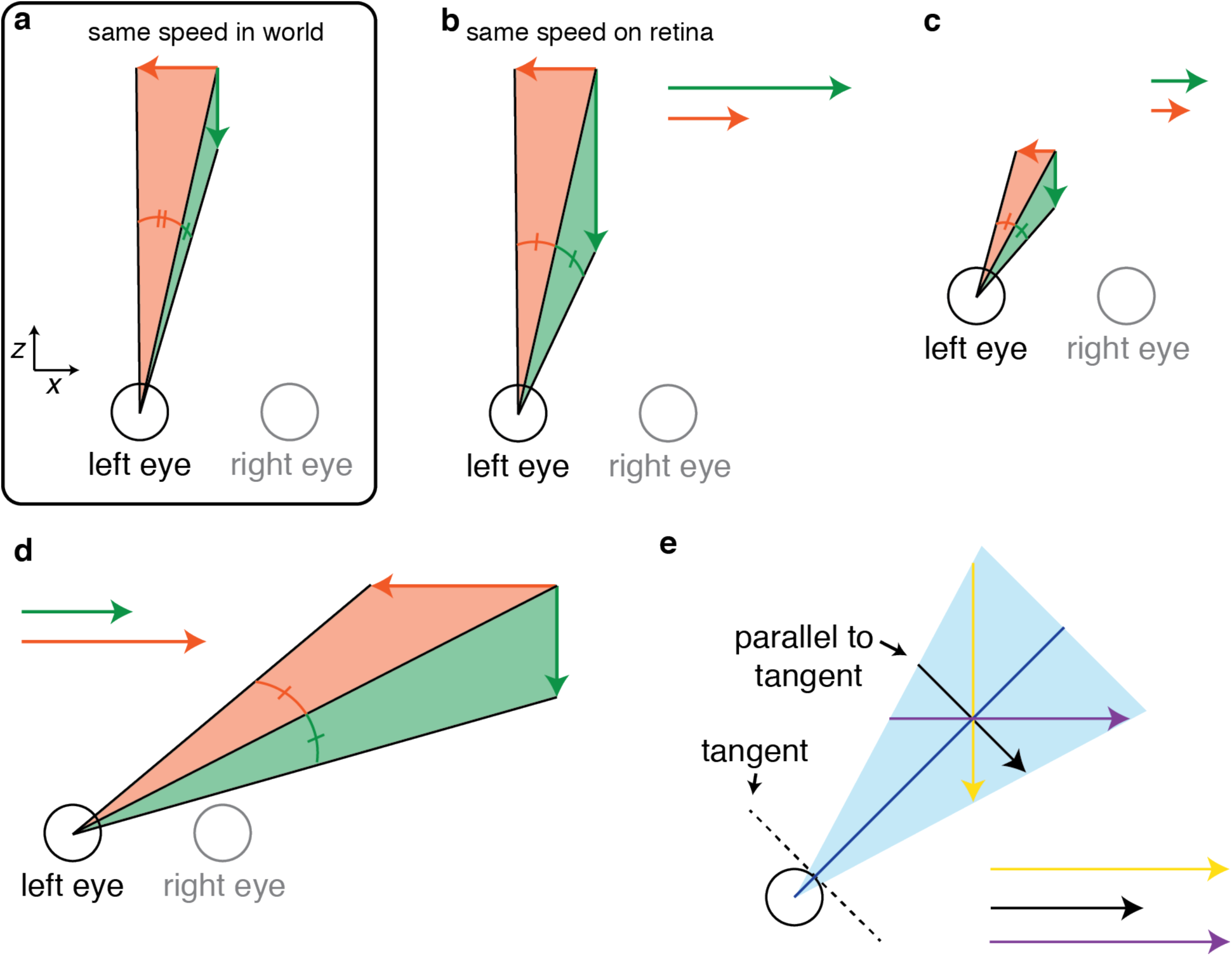
Schematic top down view illustrating how uncertainty in retinal velocity propagates asymmetrically to motion trajectories in the world. **(a)** Two orthogonal motion vectors with the same speed in the world (motion-in-depth in green and lateral motion in orange) project to different angular speeds on the retina. **(b)** A fixed retinal speed projects to a larger speed for motion-in-depth than for lateral motion. The same geometry applies to the transformation of uncertainty. **(c)** This difference is much reduced at near viewing distances. **(d)** This relationship can invert for trajectories that occur off of the midline. **(e)** Illustration of how the tangent line of a circle determines the vector direction with the minimum length for a given angle and distance. Note that when motion is directly towards either eye, this will project to zero retinal velocity in one eye (ignoring looming/optical expansion) and non-zero velocity in the other.

The angular subtense of each vector on the left eye is illustrated by the green and orange arcs, respectively. Note that although the vectors have the same length, and thus the same world speed, the angular subtense of the vector corresponding to motion-in-depth is considerably smaller than the one corresponding to lateral motion, and thus produces a considerably smaller retinal speed.

Next, we consider that our perception of motion in the world (i.e., the motion vectors) relies on measuring these angular speeds (i.e., the arcs) and inferring the physical motion trajectories that caused them. To examine how the limits of the ability of the visual system to accurately encode angular speed on the retina propagate to different motion trajectories, we can project a fixed uncertainty in angular speed back into the world. This is illustrated in **Figure 2b**, again just for two sample directions of motion directly towards or to the left of the observer. Note that even though the angular speeds are the same, the amount of uncertainty for motion-in-depth (represented by the vector length) is greater than for lateral motion. That is, a given uncertainty in angular speed results in greater uncertainty for motion-in-depth in the world. Vectors are reproduced side-by-side on the right for clarity. This difference is simply due to inverting the projection shown in **Figure 2a**. Based on this observation, it is clear that the motion-in-depth stimulus will have greater sensory uncertainty for speed estimation in the world.

However, is the high uncertainty for motion-in-depth universally true for all viewing situations? Simple geometric diagrams show that this is not the case. **Figures 2c** and **d** illustrate two additional situations. In **Figure 2c**, the distance of the motion vectors from the eyes is decreased. Uncertainty is still larger for motion-in-depth, but the increase relative to lateral motion is substantially attenuated. In **Figure 2d**, the motion vectors are located off to the observer’s right. In this case, the relationship has actually inverted and uncertainty for lateral motion is greater. Note that we only illustrate motion directly lateral to or towards the observer. However, as we will show below, since any motion vector can be decomposed into its components along these orthogonal axes, these general principles will hold for any direction.

Indeed, if we model the eye as a circle and assume the center of projection is at the circle’s center, it is easy to see that there is no consistent increase in uncertainty for motion-in-depth relative to lateral motion at all. Intuitively, for a given distance, whichever vector is closest to being parallel to the tangent of the circle (at the point where a line connecting the center of projection to the vector intersects the circle) will have the least uncertainty (**Figure 2e**). In the derivation that follows, we quantitatively determine the predicted uncertainty for 3D motion trajectories in all directions under any viewing situation and use these predictions to formulate a Bayesian model for 3D motion perception.

### Relationship between 3D motion trajectories and retinal velocities

We can describe the motion of any object in space relative to an observer in a 3D coordinate system with the function

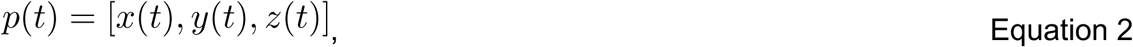

where *p* is the position of the object as a function of time (*t*) in a coordinate system defined over *x*, *y*, and *z* axes. Here, we use a head-centered coordinate system and place the origin at the midpoint between the two eyes of the observer (see icon in upper left corner of **Figure 3**). In this left-handed coordinate system, the *x*-axis is parallel to the inter-ocular axis (positive rightward), the *y*-axis is orthogonal to the x-axis in the plane of the forehead (positive upward), and the *z*-axis extends in front and behind the observer (positive in front).

**Figure 3.**
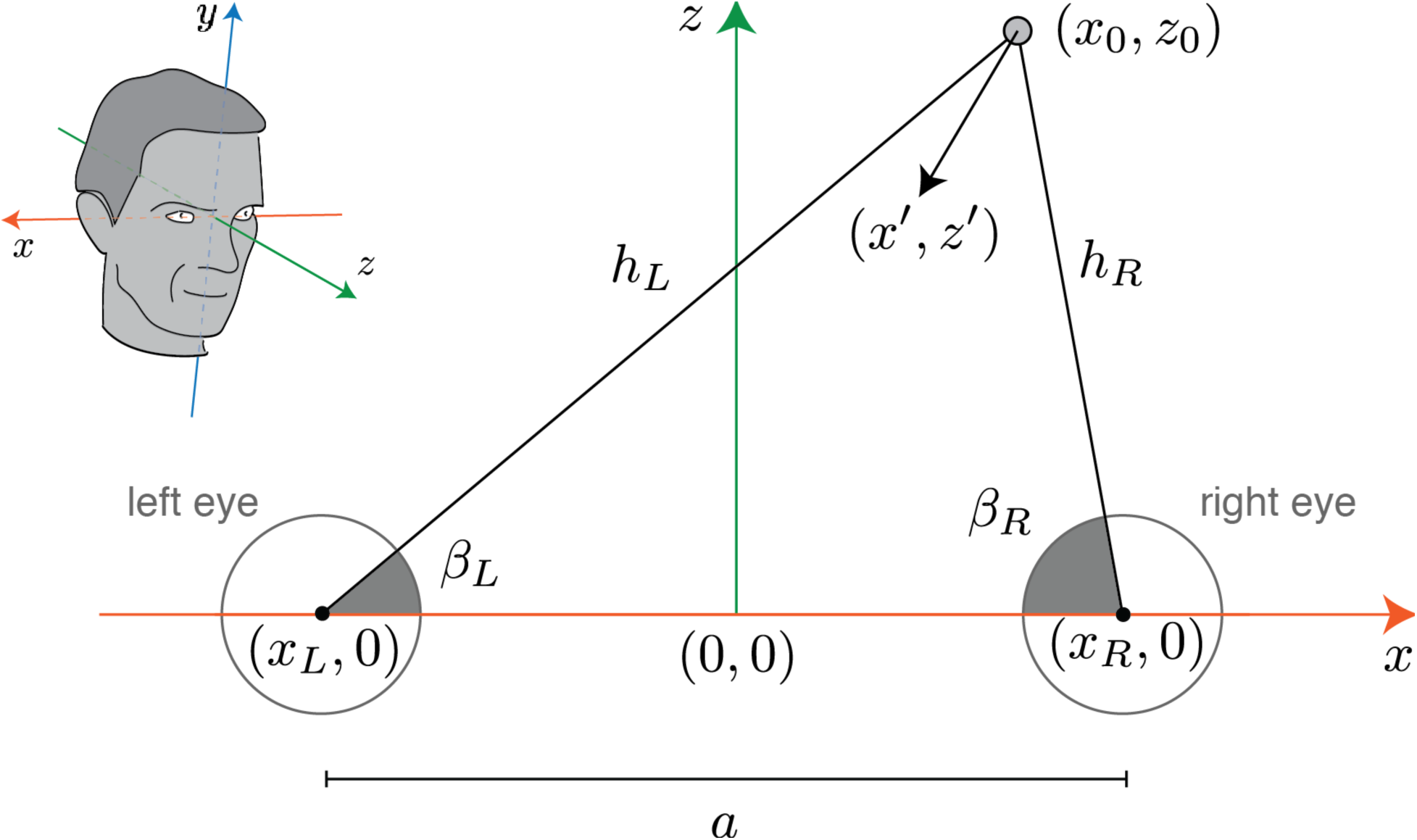
Diagram of 3D motion coordinate system. The icon in the upper left shows the origin and axes of the coordinate system, with arrowheads indicating the positive direction on each axis. The top-down view shows a slice through the inter-ocular axis in the *xz*-plane. Large circles indicate the left and right eyes. The smaller gray circle and arrow indicate the location and trajectory of motion of an object. The coordinates of key points are indicated in *x* and *z* (*y*=0 for all points), as well as several line segments and angles. Note that *x_0_* and *z_0_* denote the coordinates of the object with the motion defined by Equation 2, evaluated at time point *t*=*t_0_*.

We will model the retinal information available from horizontal velocities, and thus consider the projection of points onto the *xz*-plane (*y*=0 for all points) (**Figure 3**). Note, however, that this does not mean that this model is only valid for stimuli in the plane of the interocular axis. As long as retinal angles are represented in an azimuth-longitude coordinate system, the horizontal retinal velocities can be computed from the *x* and *z* components of 3D motion vectors alone. Also note that this geometry is independent of the point of fixation but assumes that fixation does not change over the course of the stimulus motion. In this coordinate system, the (*x*,*z*) coordinates of the left and right eye are defined as (*x_L_*,0) and (*x_R_*,0), respectively. The distance between the eyes along the inter-ocular axis, denoted by *a*, is *x_R_* – *x_L_*.

At any time point, an object with coordinates (*x*(*t*), *z*(*t*)) will project to a different horizontal angle in each eye. If we define these angles relative to the x-axis in the *xz*-plane, they are:

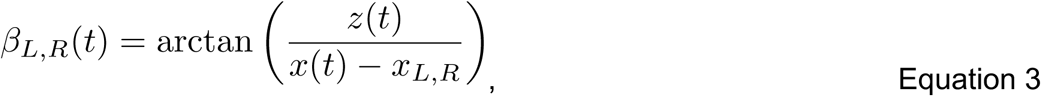

where *β_L_*(*t*) and *β_R_*(*t*) indicate the angles in the left and right eye, respectively. The object will generally have a different distance from each eye. These distances are given by:

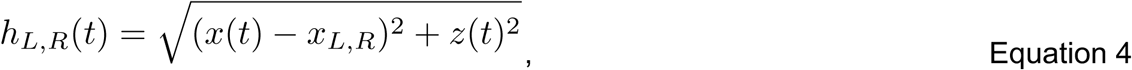

where *h_L_(t)* and *h_R_(t)* indicate the distance from the left and right eye, respectively.

Since we are interested in motion cues, we differentiate Equation 3 with respect to time to determine the relationship between object motion and motion on the retina. Here, we denote first derivatives of functions with the convention *df*(*x*)/*dt* = *f’*(*x*). This yields:

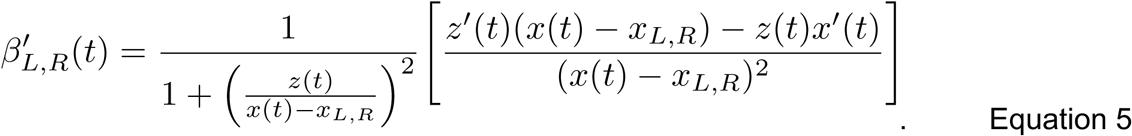

Rearranging Equation 5 and substituting in Equation 4 allows us to simplify to:

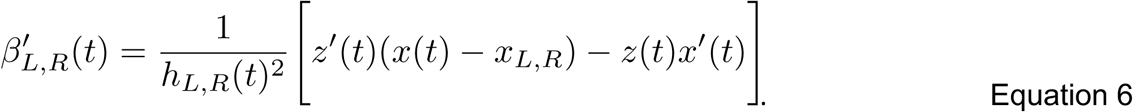

In the case of motion estimation, *β_L_*’(*t*) and *β_R_*’(*t*) are the sensory signals, and the object motion in the world that generated them are unknown. We next solve for *x*’(*t*) and *z*’(*t*) as a function of *β_L_*’(*t*) and *β_R_*’(*t*). From now on, we will define *β’_L,R_*, *h_L,R_*, *z_0_*, *z’*, *x_0_*, and *x’* to be equal to *β’_L,R_*(*t*), *h_L,R_*(*t*), *z*(*t*), *z*’(*t*), *x*(*t*), and *x*’(*t*), each evaluated at the same time point *t=t_0_*. To determine the equation for *x*’ in terms of retinal velocities, we rearrange Equation 6 for the left eye to solve for *z*’, substitute the result back into Equation 6 for the right eye, and solve for *x*’, yielding:

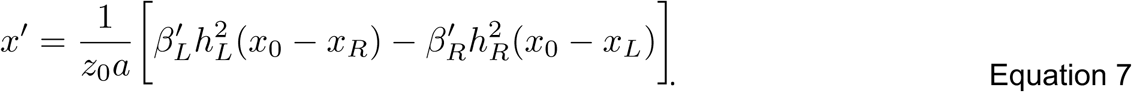

Recall that *a* refers to the interocular separation. To determine the equation for *z*’ in terms of retinal velocities, we rearrange Equation 6 for the left eye to solve for *x*’, and substitute this back into Equation 6 for the right eye, yielding the equation for *z*’, also in terms of retinal velocities:

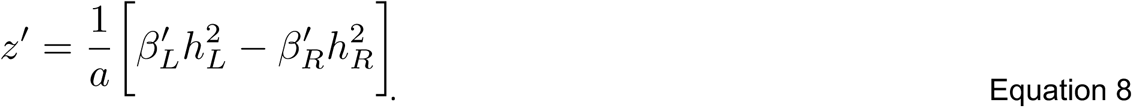

### Propagating uncertainty for 3D motion

We assume that the measurements of retinal motion in each eye, *β’_L_* and *β’_R_* are corrupted by independent additive noise:

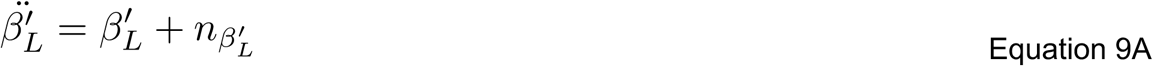

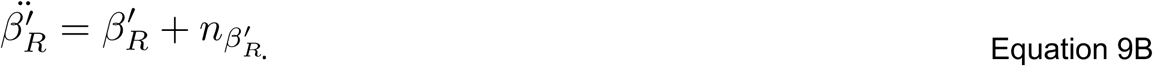

Here, 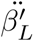 and 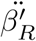 denote the measured retinal velocities in the left and right eye, respectively, and noise samples (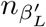 and 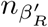) are drawn independently for both eyes from a zero mean Gaussian distribution with variance of 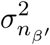. Note that the assumption of constant additive noise is inconsistent with Weber’s Law (which would predict that the noise increases proportionately with speed). However, psychophysical experiments have shown that for relatively slow speeds (less than ~2-4 deg/s), speed discrimination thresholds are more stable than predicted by Weber’s Law (McKee, Silverman, & Nakayama, 1986; Stocker & Simoncelli, 2006; Freeman, Champion, & Warren, 2010). Most stimuli in our experiments moved at speeds slower than 4 deg/s.

We should note that our derivation only depends on the locations of the eyes and the object, and is independent of where the observer fixates. Under the assumption that the object’s 3D location – its distance *z_0_* and its location relative to each eye (*x_0_* − *x_L_*) and (*x_0_* − *x_R_*) – are known, we can use Equations 7 and 8 (which specify *x’* and *z’* as linear combinations of 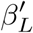 and 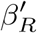) to determine the noise covariance of the sensory measurements of speed in *x* and *Z*(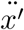 and 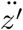). First, we rewrite the linear transformation from retinal velocities 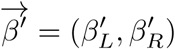 to real-world velocities 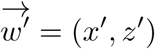 in terms of the matrix equation 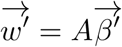. In this formulation, A is given by:

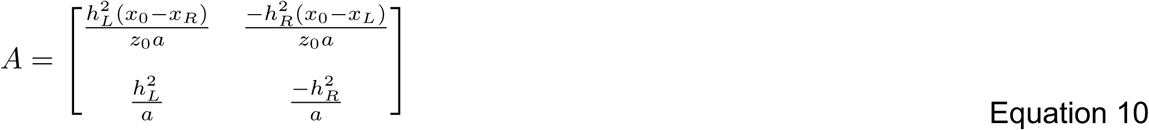

If we assume independent and equal noise distributions for the two eyes, the noise covariance of the sensory measurements (denoted *M*) is given by 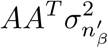:

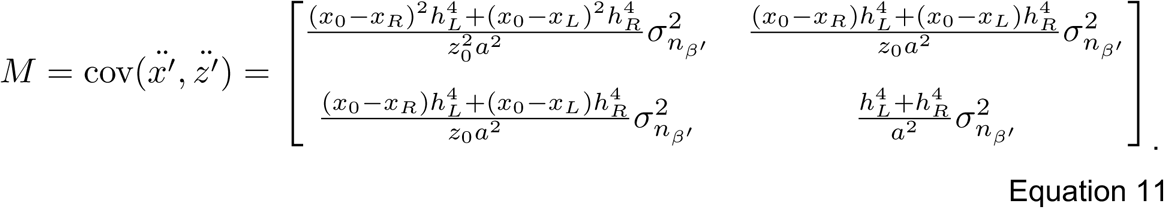

To gain more intuition for the relative noise effects on the *x* and *z* velocity components of a motion trajectory, we plot the sensory uncertainty (the noise standard deviation) for each velocity component (the square root of the diagonal elements of Equation 11, denoted *σ_x′_* and *σ_z′_*) as a function of horizontal distance (*x_0_*) and distance in depth (*z_0_*) in **Figure 4a** and **b** (assuming an interocular distance *a* = 6.4cm). Each panel contains an isocontour plot showing the log of the sensory uncertainty at each true spatial location. Several features are notable. The uncertainty in *x’* is at its minimum for points that fall on or near the midsagittal plane, and increases for points to the left and right. The uncertainty in *z’* is at its minimum for points closest to the eyes, and increases radially away from the mid-point between the eyes. Note that uncertainty in *x’* also increases with distance, but not as sharply as *z’*. In the central visual field, the uncertainty in *z’* is generally much greater than the uncertainty in *x’*.

**Figure 4.**
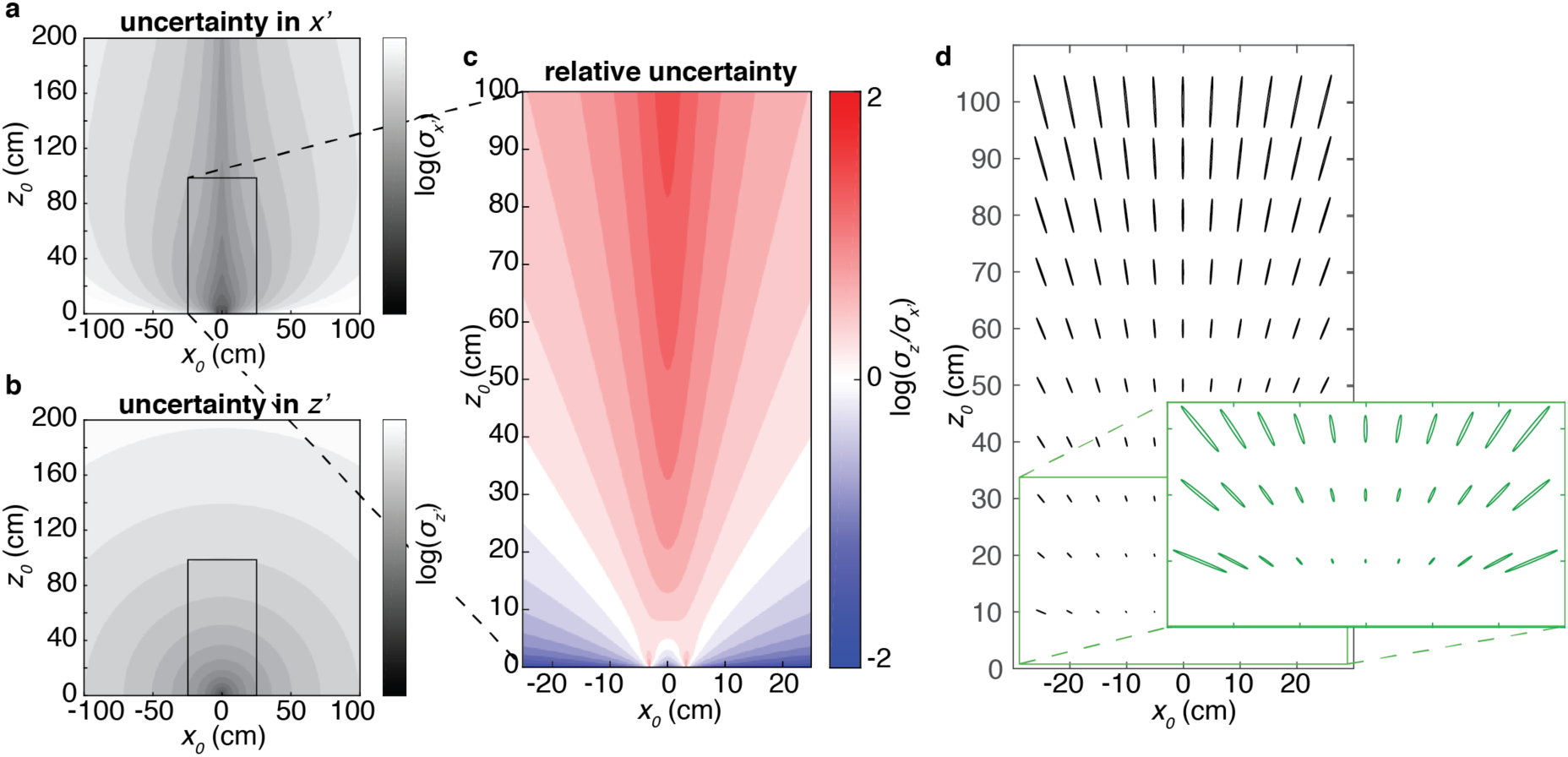
Uncertainty for *x* and *z* motion vary with stimulus distance and head-centric eccentricity. **(a)** Uncertainty in the *x* component of a motion vector is plotted in arbitrary units as a function of location in *x* and *z*. **(b)** Same as (a), except for the *z* component of motion. The color map scales of panels a and b are the same. **(c)** The ratio between the values in the boxed region in (a) and (b). **(d)** Ellipses illustrate the noise covariance of *x’* and *z’* for a range of spatial locations. Ellipse scale indicates the relative uncertainty for each location, and orientation indicates the axis of maximal uncertainty. All ellipses have been reduced by scaling with an arbitrary factor to fit within the plot. Inset shows the same ellipses for a small spatial region (also with a different scaling).

To illustrate the relative magnitude of uncertainty in *x’* and *z’*, we plot the ratio of the two values for a subset of points close to the observer in **Figure 4c** (within 25cm left/right and 100cm in depth). Ratios greater than 1 (red) indicate that uncertainty in *z’* is greater than *x’*, and ratios less than 1 (blue) indicate the reverse. In the central visual field, this ratio is greater than 1. This is consistent with previous work (Welchman et al., 2008). However, the ratio varies considerably as a function of both viewing distance and viewing angle. At steep viewing angles (> 45 degrees), the relationship reverses and *x’* uncertainty is actually greater than *z’*. We should note that our model only includes uncertainty in object speed, not in object location. Uncertainty in object location would likely increase for objects projecting to larger retinal eccentricities.

Equation 11 indicates that the uncertainties in *x’* and *z’* are not independent. To visualize this relationship, in **Figure 4d**, we show the covariance ellipses for a set of locations within 100cm in depth (the inset shows a zoomed view of the points with 50cm in depth). For most locations, the ellipses are highly elongated, indicating that for each location, uncertainty about different motion components differs strongly. As expected from the geometric analysis (**Figure 1**), the axis of minimal uncertainty is orthogonal to a line connecting each location back to the interocular axis, independent of the direction of gaze. This creates a radial pattern, in which uncertainty is highest for motion extending radially from the observer’s location. Along the midsagittal plane (*x_0_* = 0), the covariance is zero and the axes of minimal and maximal uncertainty align with the x and z axes, respectively.

Indeed, if we consider only cases in which the stimulus is presented in the midsagittal plane, as is often done in perceptual studies, the off-diagonal elements of the covariance matrix become zero, and we can simplify 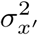 and 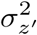 to:

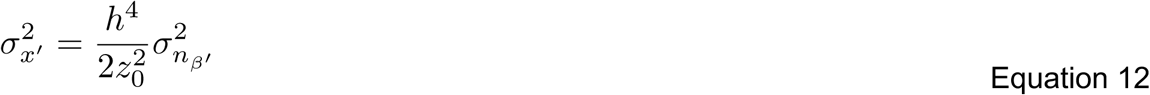

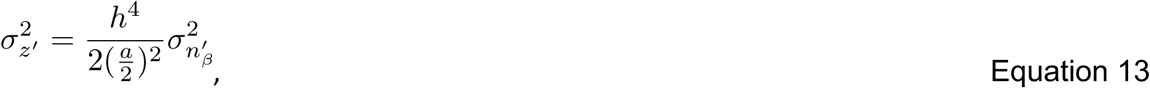

where 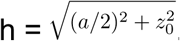. During typical viewing *a* << *z_0_*, resulting in substantially larger uncertainty for the *z* component of velocity than for the *x* component. However, if *z_0_* is equal to half of the interocular separation, the variances in *x’* and *z’* will be equal as well. Thus, while uncertainty for motion in depth in the midsagittal plane clearly tends to be substantially higher than for lateral motion, the relative uncertainty is reduced for near viewing distances.

### Application of Bayes rule to predict perceived motion in the midsagittal plane

In order to predict how sensory uncertainty in motion measurement affects actual percepts, we need to define a formal relationship between sensory measurements and perceived motion. For this, we derive a Bayesian ideal observer that combines this sensory information with a prior distribution over 3D motions. We will first consider only motion in the midsagittal plane (*x_0_* = 0), such that uncertainty in *x’* and *z’* from the likelihood come out as independent (Equations 12 and 13).

The full likelihood function in real-world coordinates – that is the conditional probability of the (transformed) velocity measurements 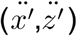 given the true velocities (x’, z’) – can be given by the a 2D Gaussian probability density function. For simplicity, we specify this as the product of two 1D Gaussians, 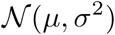, where *μ* and *σ* denote the mean and standard deviation of a 1D Gaussian, respectively. Thus, for motion in the midsagittal plane :

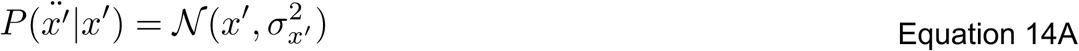

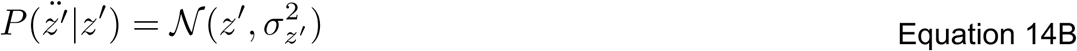

where 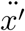 and 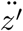 are derived from the measurements of retinal velocity in the left and right eyes (Equations 7-9).

We assumed that the prior for slow speeds is isotropic in world coordinates (a Gaussian with equal variance in all directions) and centered at zero, as has been done previously (Welchman et al., 2008, but see Lages, 2006 for an alternative approach). We can then express the prior as:

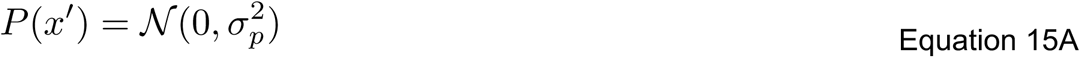

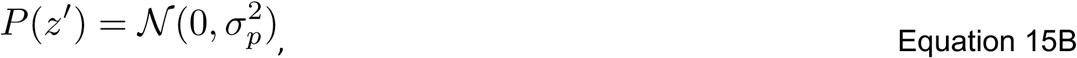

where 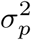 is the variance of the prior.

The posterior distribution, according to Bayes’ rule, results from multiplying the likelihood and prior, and renormalizing. For Gaussian likelihoods and priors, the posterior distribution also takes the form of a Gaussian, with mean and variance that can be computed according to standard formulas. The means of the posterior in x’ and z’ are given by:

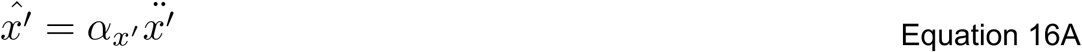

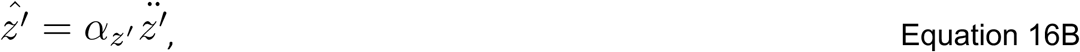

and the variances by:

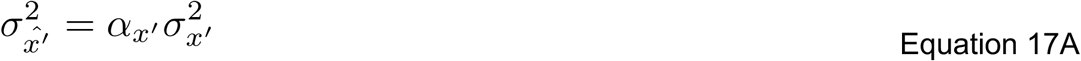

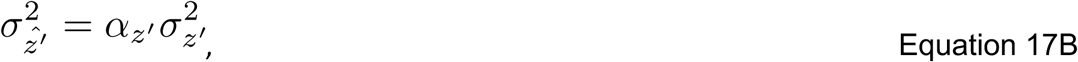

where these means are denoted by 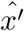 and 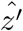 because they also correspond to the sensory estimate of each motion component determined by the maximum a posteriori (MAP) method. In Equations 16 and 17, *α* is a “shrinkage factor” that determines how much the measured velocity components are shrunk towards zero (the mean of the prior). The equations for each factor (*α_x′_* and *α_z′_*) are:

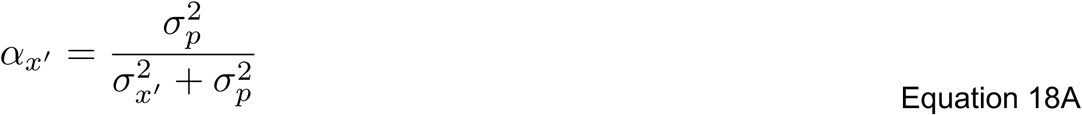

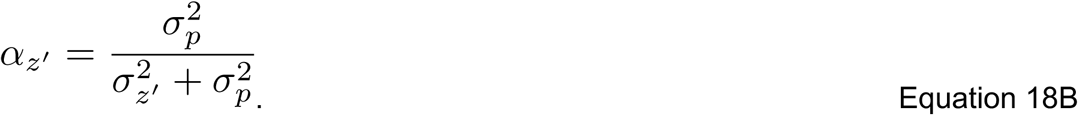

In brief, the estimated speeds correspond to the measured speeds in each direction, scaled towards zero by the shrinkage factor in each direction. Similarly, the posterior variance equals the variance of the sensory measurements also scaled by the shrinkage factor.

The full posterior distribution, that is, the probability of a given world velocity given a particular measured velocity, can therefore be written:

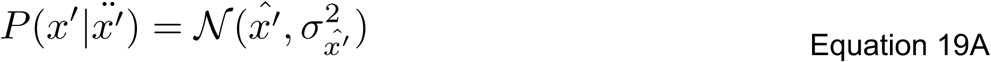

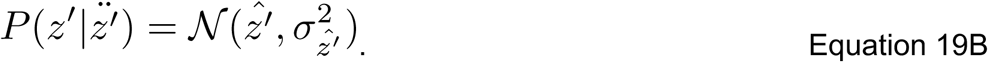

We can examine the trial-to-trial performance of the Bayesian ideal observer by deriving the sampling distribution of the MAP estimate, that is, the distribution over the estimates of a Bayesian ideal observer given a fixed stimulus. (The ideal observer exhibits variability because it receives a new set of noisy measurements on each trial). This distribution is given by:

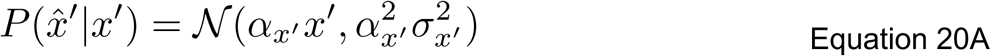

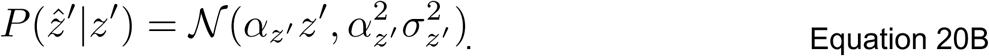

Here *α_x′_* and *α_z′_* can again be understood as “shrinkage factors” that determine how much the observer’s reports are scaled towards zero (on average) relative to the true velocity. The variance of the ideal observer’s estimates are scaled by 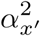 and 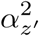 relative to the variance of a maximum likelihood estimator (due to the fact that a Gaussian random variable scaled by *α* will have its variance scaled by *α*^2^). This shows that the ideal observer exhibits a reduction in variance even as it exhibits an increase in bias (in this case, bias towards slower speeds).

### Application of Bayes rule to predict perceived motion at arbitrary locations

For the case of motion occurring away from the midsagittal plane, we can derive the full covariance matrix of the ideal observer’s estimates (which is not aligned with cardinal x/z axes). We already have the noise covariance of the sensory measurements from Equation 11. The covariance of the posterior of the Bayesian ideal observer (denoted by Λ, the covariance of 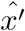 and 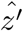) can be determined from this matrix (*M*) and the covariance of the prior (denoted as *C*, a diagonal matrix with variance in *x’* and *z’* of 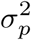:

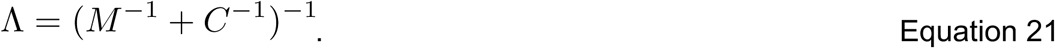

Given a pair of sensory measurements 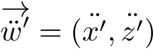, the vector of posterior means in *x’* and *z’* (i.e., 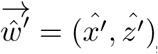, the MAP estimate in *x’* and *z’*) is then:

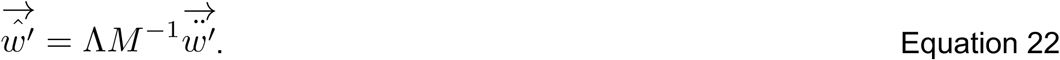

Here, the matrix *A* = Λ*M*^−1^ provides a “shrinkage factor” on the maximum likelihood estimate analogous to the role played by *α* in the previous section.

Lastly, the sampling distribution of the MAP estimate can be described as a 2D Gaussian:

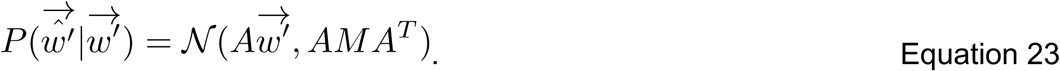

In the next sections, we describe the results of our 3D motion perception experiments and compare the results to the predictions of this ideal observer model. For all model predictions, we assumed an observer with a typical interocular separation of 6.4 cm and a stimulus speed of 1 cm/s (typical of the experiment). Model predictions for a range of speeds presented in the experiment were qualitatively similar. With the mean of the likelihood determined by the stimulus speed and the viewing geometry, the variance of the sensory noise is not specified by the model, and neither are the mean and variance of the prior. To address this, the mean and standard deviation of the prior were fixed at 0 and 1 cm/s, respectively. Differences in stimulus contrast were modeled as differences in the variance of the retinal velocity measurement noise (which determines the variance of the likelihood). This variance was selected such that the Bayesian model predicted a lateral angle bias that matched the average bias for one condition in Experiment 1. Specifically, we matched the prediction for the stimulus with 100% contrast at 45cm, the condition with the least perceptual bias. All other predictions were then made based on this parameter, assuming that reductions in stimulus contrast resulted in a proportionate increase in the noise variance. Since the Bayesian model considers only one 3D motion cue, and the experimental stimuli contained a range of cues, analyses were performed to explore whether the overall patterns predicted by the model were borne out in the data, not with the goal of numerically fitting the model to the experimental data.

We summarized both model and experimental data using various descriptive statistics. First, we examined the *average* errors predicted by the Bayesian model for each direction of motion in the world. This is simply the derived MAP for each possible motion direction between 0 deg and 359 deg in steps of 1 deg (**Figure 1c**). Next, to examine the trial to trial variance of the Bayesian model (and therefore determine the predicted percentage of motion direction misperceptions), we examined the sampling distribution of the MAP and determined the percentage of MAP samples that would result in judgments in the opposite direction of the stimulus for motion-in-depth (towards versus away) and lateral motion (leftward versus rightward). Model predictions were otherwise analyzed using the same methods described earlier for the experimental data.

### Predicted and observed biases towards lateral motion in the midsagittal plane vary with stimulus distance and contrast

Recall that previous perceptual experiments have demonstrated that observers tend to overestimate the angle of approach of objects. That is, an object on a trajectory towards the observer tends to be perceived as moving more laterally than the true stimulus trajectory. We refer to this as a ‘lateral bias.’ **Figure 5a** clearly shows that the current model predicts this lateral bias. Here, the simulated stimulus is located at 90cm directly in front of the observer (i.e., *x_0_* = 0, *z_0_* = 90). For simplicity, we convert the stimulus motion (both actual and predicted) into a single direction value in degrees, given by the angle between the motion trajectory and a vector centered on the object and parallel to the x-axis in the positive direction (i.e., 0 deg = rightward motion, see inset). Counterclockwise angles are positive. The direction of stimulus motion (*θ*) was varied across the full 360 deg range (abscissa) and the predicted perceived direction for each value was calculated (ordinate). Arrows indicate motion directions from a top-down view as shown in **Figure 1**. Veridical perception is indicated by the dashed diagonal line. The weaving pattern of the predictions indicates that on average the motion-in-depth trajectories, except for those that are directly towards or away, tend to get pulled towards more lateral motion. For example, when the stimulus moves directly towards the observer (270 deg), the average predicted percept is veridical (also 270 deg). However, when the trajectory towards the observer is slightly off to the left or right, the average predicted percept is biased more leftwards or rightwards. The same is true for trajectories away from the observer.

**Figure 5.**
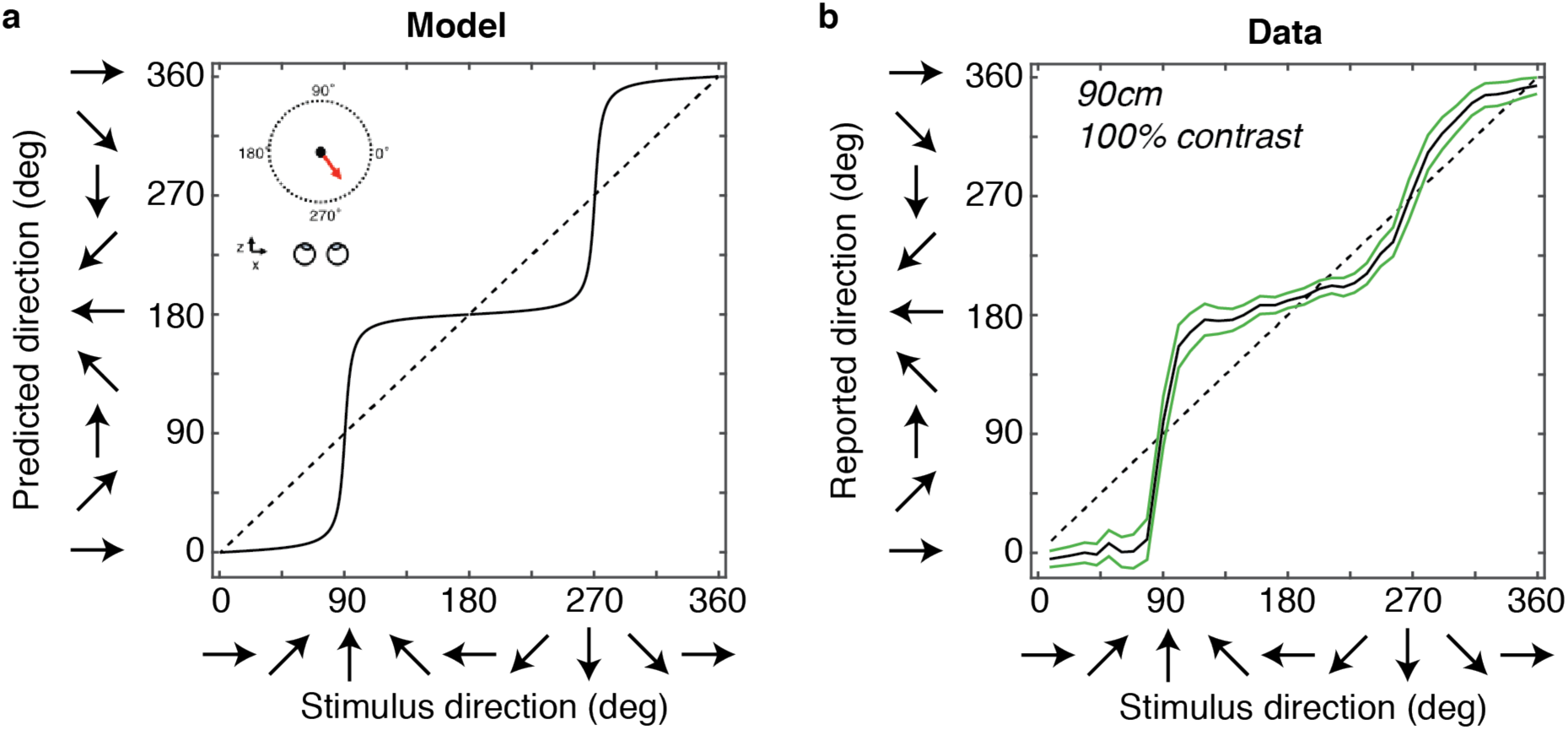
Comparison between model predictions and average human performance in Experiment 1. **(a)** Predicted reports of perceived target direction as a function of the true stimulus direction for a participant viewing a stimulus from a distance of 90 cm. **(b)** Averaged results for the 15 participants who took part in the experiment viewing the stimulus within the virtual-reality environment at 90 cm in the same format as (a). Data are binned in 10 degree increments. Error bars correspond to +/-1 standard error of the mean (SEM) across participants.

However, these model predictions are made based only on retinal motion cues, and during natural behavior there are many other cues available that may improve the estimate of an object’s motion trajectory. To determine the extent to which these biases contribute to 3D motion perception during natural behavior, we conducted an experiment in a virtual environment (Experiment 1). The stimulus was rendered with perspective projection, so that monocular cues (looming) were available. In addition, participants were free to move their head during the experiment, and the view of the stimulus was updated based on any head rotation. Thus, the use of the virtual reality headset made some extra-retinal head motion cues available as well. For comparison, in **Figure 5b** we show the average perceived direction for the same range of motion trajectories presented to observers in this virtual reality environment (90 cm distance, 100% contrast). The weaving pattern again indicates that there is a lateral bias in these experimental data, as predicted by the model. The presence of other cues may reduce the lateral bias, but they do not eliminate it.

**Figure 6** expands upon this result and shows how this bias changes as a function of two stimulus features: the viewing distance and the contrast of the stimulus. Each bar indicates the average signed error between the stimulus and the percept predicted by the model (**Figure 6a**) or the measured participant responses from the experiment (**Figure 6b**). Larger values of this error indicate larger laterality biases (see Methods). The model (**Figure 6a**) predicts a decrease in the laterality bias with decreased viewing distances (i.e., 45 cm versus 90 cm). The model also predicts an increase in the laterality bias for lower contrast stimuli. This predicted increase is more substantial for the near viewing distance (dark bars), and is relatively weak for the farther viewing distance (light bars), for which the lateral bias is substantial at all contrasts. Both of these effects are present in the observed errors in the behavioral experiment (**Figure 6b**); however, the overall magnitude of the errors is smaller. Because the model considers only one depth cue (motion), and the experiments contain many more, it is reasonable to expect a somewhat reduced magnitude of the observed errors. Recall that the model was matched to the data using a single free parameter: setting the variance of the likelihood for a single stimulus (45cm distance, 100% contrast).

**Figure 6.**
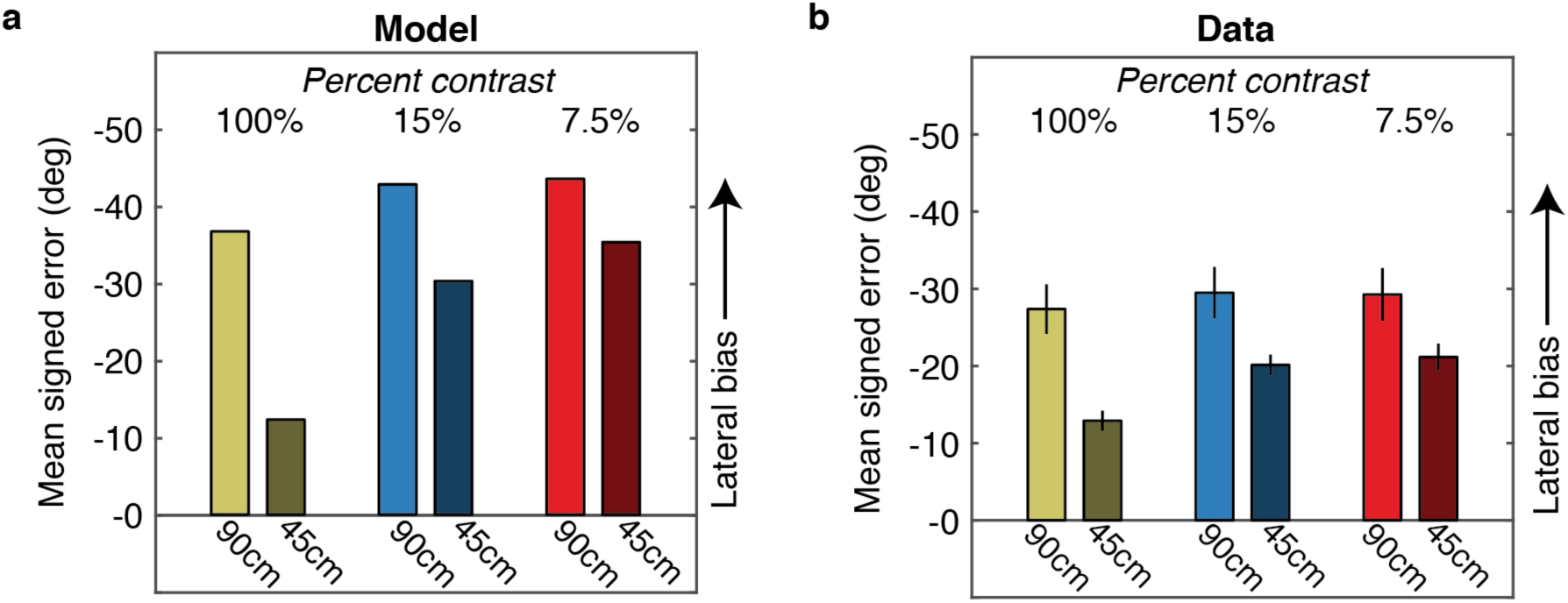
Comparison between model predictions and human lateral bias in Experiment 1. **(a)** Mean signed error in predicted perceived target direction for viewing three target contrast levels at two viewing distances. Negative values (increasing on the ordinate) correspond to reports that are laterally biased. **(b)** Results for the 47 participants (n=15 for 90 cm and n=32 for 45 cm) who took part in the experiment viewing the stimulus within the virtual-reality environment in the same format as (a). Error bars correspond to +/-1 SEM.

A two-way ANOVA performed on the experimental data showed a significant main effect of viewing distance on human performance (*F(1,135)=6.67, p=0.01*), with a reduction in perceptual bias for object motion nearer to the head. Moreover, the data reveal the predicted effect of target contrast on human performance. Specifically, viewing distance interacts with target contrast. Multiple comparisons revealed that the relative decrease in perceptual bias at the nearer viewing distance is significantly greater for the mid and high target contrast levels *(p<0.0167, Bonferroni-corrected alpha)*. The perceptual bias is also impacted by an increase in viewing distance for low contrast targets, however, the difference reported here failed to reach significance *(p=0.026)* at the Bonferroni-corrected alpha level.

While a previous model did predict an effect of viewing distance (Welchman et al., 2008), prior experimental studies concluded that distance does not modify the lateral bias (Harris & Dean, 2003; Poljac, Neggers, & van den Berg, 2006). Until now, this inconsistency between model and data had not had a clear explanation. However, as demonstrated in **Figure 6a**, the amount of the predicted difference between viewing distance interacts with other properties of the stimulus uncertainty (here shown as contrast, but generally summarized as the variance of the likelihood). Thus, it is possible that some experimental set ups would reveal a distance effect, and others might not, particularly with relatively small sample sizes such as those used in the previous studies (3 and 6 participants, respectively). The current set of experimental data are clearly consistent with an impact of Bayesian inference on biases in 3D motion perception.

In addition to the dependence of the laterality bias on viewing distance, the model also predicts a dependence on stimulus eccentricity. The model predicts that the relative stimulus uncertainty in depth (and therefore the laterality bias) should be reduced when an object is located off to the left or right, rather than directly in front of the observer (**Figures 2 and 4**). We will return to this prediction in last section of the Results.

### Misperceptions in motion direction in the midsagittal plane

Recent work has shown that motion trajectory judgments can be subject to direction reversals for approaching and receding motion, but much less so for leftward and rightward motion (Fulvio et al., 2015). That is, observers sometimes report that approaching stimuli appear to move away, and vice versa, but rarely if ever report leftward-moving stimuli as appearing to move rightward, and vice versa. Can such reversals be explained by our Bayesian model? The preceding analysis of laterality biases is based on taking the MAP in order to predict the average perceived motion trajectory for a given stimulus. However, the Bayes observer model is fundamentally probabilistic. On any given stimulus presentation, the percept is expected to be drawn from the sampling distribution of the MAP.

To examine whether perceived direction reversals can be accounted for by the model, we first plot the full sampling distribution of the MAP for two example stimuli: motion directly towards an observer (270 deg) and motion directly to the right (0 deg) (the left panels of **Figure 7a** and **b**, respectively). Specifically, for each example stimulus, we show a heat map of this distribution, with *x’* plotted on the horizontal axis and *z’* plotted on the vertical axis. These plots demonstrate that a large percentage of the sampling distribution for a stimulus moving towards an observer can occur for trajectories that recede in depth. In other words, the variance of the MAP sampling distribution in the *z*-direction can be large enough so that it extends into the opposite direction of motion. For rightward motion however, very little of the distribution occurs for leftward trajectories, and vice versa. To examine the percentage of trials in which observers should misreport motion direction, we converted the trajectories in the sampling distribution of the MAP to direction angles and replotted the normalized frequency in polar coordinates as a function of motion direction (**Figure 7a** and **b**, right panels). Non-zero values in the opposite direction of motion (away or leftward) indicate that the model predicts a certain percentage of trials will include direction confusions.

**Figure 7.**
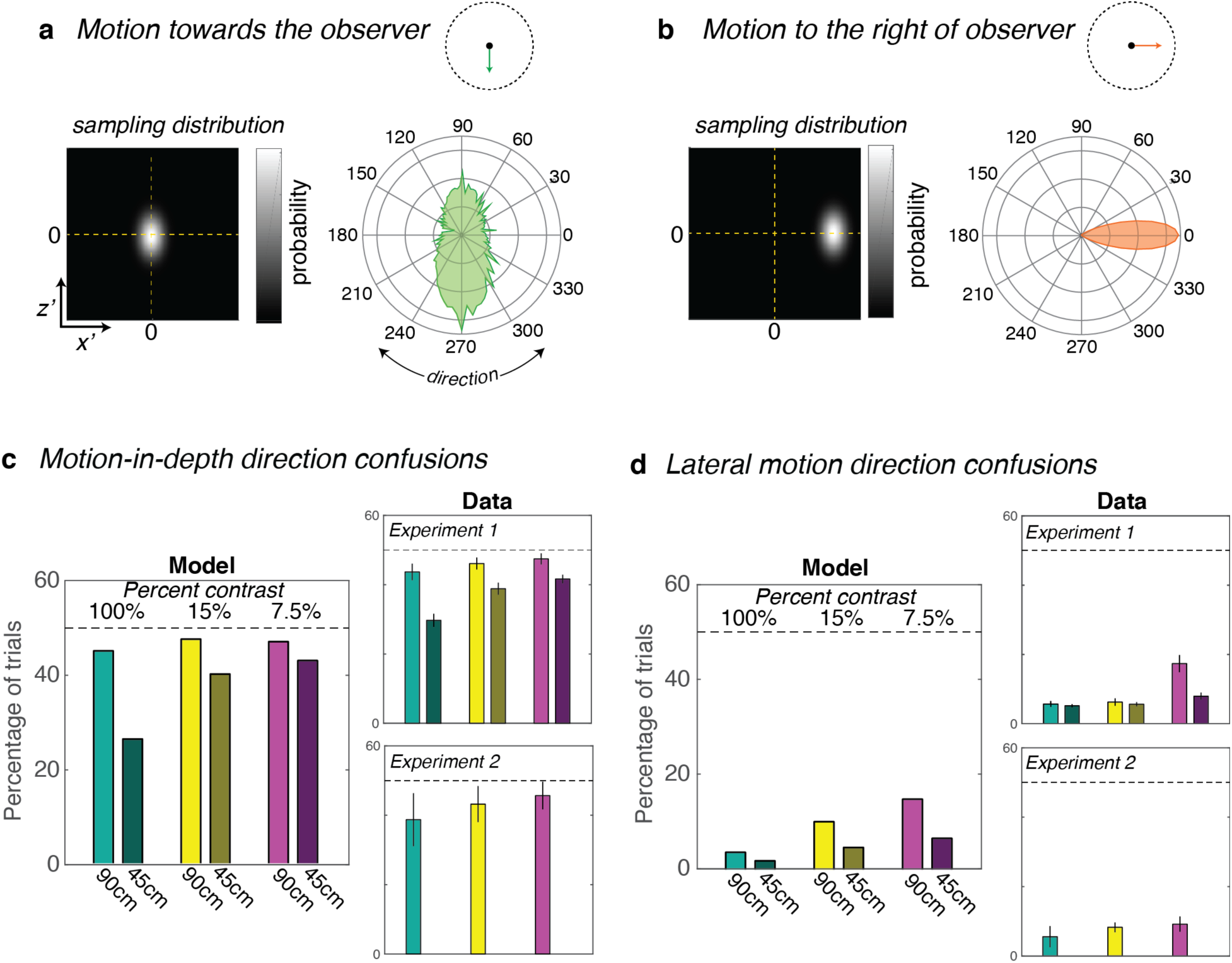
Direction confusions for motion-in-depth and lateral motion. **(a,b)** Illustrations of the predicted sampling distribution of the MAPs for motion directly towards an observer (a) and directly to the right of an observer (b) in Cartesian and polar coordinates. Model parameters used were for a 90cm viewing distance, and 100% stimulus contrast. **(c,d)** Predictions of the model and experimental results for motion-in-depth confusions (c) and lateral motion confusions (d). Experiment 1 and Experiment 2 are shown in separate panels. Error bars correspond to +/-1 SEM.

Next, we examined the effects of distance and contrast on predicted and observed direction confusions, averaging across all directions of motion in the world. **Figure 7c** and **d** show that the Bayesian model predicts that direction confusions for motion-in-depth (**Figure 7c**) will greatly exceed lateral motion confusions (**Figure 7d**). Each bar represents the predicted percentage of trials in which direction will be confused, and the dashed line indicates chance performance (50%).

For motion-in-depth confusions, the model predicts that direction confusions will decrease with reduced viewing distance (90 cm versus 45 cm). The model also predicts that direction confusions will increase as sensory uncertainty increases (contrast decreases from 100% to 15% to 7.5%), most markedly at the nearer viewing distance (dark bars). The upper right-hand panel in **Figure 7c** shows the results from Experiment 1, plotted in the same manner as the model predictions. Overall fewer confusions occur in the experimental data than are predicted by the model, likely due to the presence of visual cues to motion-in-depth in the experiments, that were not incorporated into the model. Importantly, however, the overall effects of stimulus distance and contrast are well-matched to the model predictions.

A two-way ANOVA conducted on the data from Experiment 1 revealed a main effect of viewing distance on human performance (*F(1,135)=5.8, p=0.02*), with a reduction in direction confusions for object motion nearer to the head. There was also a significant interaction between viewing distance and target contrast (*F(2,135)=4.3, p=0.02*). Multiple comparisons revealed that direction confusions significantly increased for all target contrast levels *(p<0.0167, Bonferroni-corrected alpha)* as the viewing distance doubled from 45cm to 90cm.

Because direction confusions might seem surprising, we compared these results to a second experiment (Experiment 2, lower right-hand panel for **Figure 7c**). This experiment used a standard stereoscopic display and a random dot stimulus. Note that Experiment 2 included a contrast manipulation, but stimuli were always presented at one distance, and the high contrast condition was 60% rather than 100% Weber contrast. As predicted, a one-way ANOVA on the data from Experiment 2 revealed a main effect of target contrast (*F(2,4)*=160.99, *p*<0.001).

The model predicts that lateral motion direction confusions will be much less frequent, but will be similarly affected by viewing distance and stimulus contrast (**Figure 7d,** left panels). That is, in the fronto-parallel plane, direction confusions will decrease with reductions in viewing distance and increase with reductions in stimulus contrast. These predicted effects were present in both experiments (**Figure 7d**, right panels). A two-way ANOVA on the data from Experiment 1 revealed main effects of both viewing distance (*F(1,135)=*32.8, *p<*0.001) and contrast (*F(2,135)*=35.2, *p<*0.001). The interaction between viewing distance and contrast was also statistically significant (*F(2,135)*=14.4, *p*<0.001). Follow up comparisons revealed that direction confusions significantly increased with viewing distance for the lowest contrast stimulus. Although the average percentage of lateral misperceptions was highest in the low contrast condition of Experiment 2, the effect of stimulus contrast was not statistically significant (*F(2,6)=3.4*, *p*=0.1).

To summarize, while overt confusions in the direction of motion seem surprising on their own, they are clearly predicted by the same Bayesian motion perception model that accounts for other perceptual phenomena.

In addition to these overall motion-in-depth direction confusions, it is well documented that motion trajectories towards the observer have some amount of ‘privileged’ perceptual processing (Lin et al., 2008, 2009; Schiff et al., 1962). Indeed, using similar random dot stimuli to those from Experiment 2, our prior work has shown an overall bias to perceive motion-in-depth stimuli as approaching rather than receding (Fulvio, Rosen, Rokers, 2015; Cooper, van Ginkel, & Rokers, 2016). Still under some (low contrast) stimulus conditions, this bias is reversed, and motion tends to be perceived as receding. Indeed, this same pattern was observed in the current experiments – for the low contrast stimuli, there were substantially more motion-in-depth misreports when the stimuli moved towards the observer (a ‘receding bias’), and for the high contrast stimuli, this tended to reverse (a ‘towards bias’). The current Bayesian model does not predict these asymmetries – although the sampling distribution of the MAP extends into reversed directions, the average of this distribution (the MAP itself) is always in the same direction as the stimulus. An additional prior assumption, a prior that shifts away from zero speeds for some stimuli, or attentional effects would need to be incorporated to account for these asymmetries in how direction confusions occur.

### 3D motion perception outside of the midsagittal plane

The previous sections have considered motion trajectories originating in the midsagittal plane. Of course, in the real world, stimuli need not be confined to this plane and may originate in any location relative to the observer. The model reveals that while the uncertainty for motion estimates in both *x* and *z* increases with distance from this plane, it does so at different rates (**Figure 4**). While uncertainty of the *z* motion estimate is typically much larger than the corresponding *x* motion estimate in the midsagittal plane, the relative uncertainty decreases away from that plane. In fact, at an angle of 45 deg away from that plane the relative uncertainty becomes unity, predicting unbiased estimates of motion trajectory. Beyond 45 degrees the relationship reverses, such that the model will predict a medial, rather than a lateral bias. Another way to think about this is that the axis of maximal uncertainty shifts from being aligned with the *z* axis in the midsagittal plane to becoming aligned with the *x* axis for motion originating directly to the left or right of the observer (see **Figure 2**). Because of this, estimated motion trajectories predicted by the model will differ between midsagittal and peripheral motion trajectories.

In Experiment 3, we tested whether the observed lateral bias and motion direction confusions are affected by stimulus location, in accordance with model predictions. **Figure 8a** and **c** show the model predictions for lateral bias and direction confusions for motion trajectories originating in the midsagittal plane (central) and 20 deg to the left or right (peripheral). At this eccentricity, the lateral bias is predicted to change only slightly, decreasing for peripheral targets by just 0.7 deg relative to central targets (**Figure 8a**). In contrast, the model predicts that the percentage of motion direction confusions will increase substantially for lateral motion (*x*) and decrease slightly for motion-in-depth (*z*) at peripheral locations, relative to the midsagittal location (**Figure 8c**). These model predictions are qualitatively similar to the experimental data: we observed a small decrease in lateral bias and motion-in-depth confusions at the peripheral location, and a large increase in lateral direction confusions (**Figure 8b** and **d**). It is also notable that performance was overall worse in this experiment (the lateral bias was larger than in the 45 cm midsagittal stimulus from Experiment 1). This is potentially because of the additional demand placed by asking the observers to attend to all three possible stimulus locations at the start of a trial.

**Figure 8.**
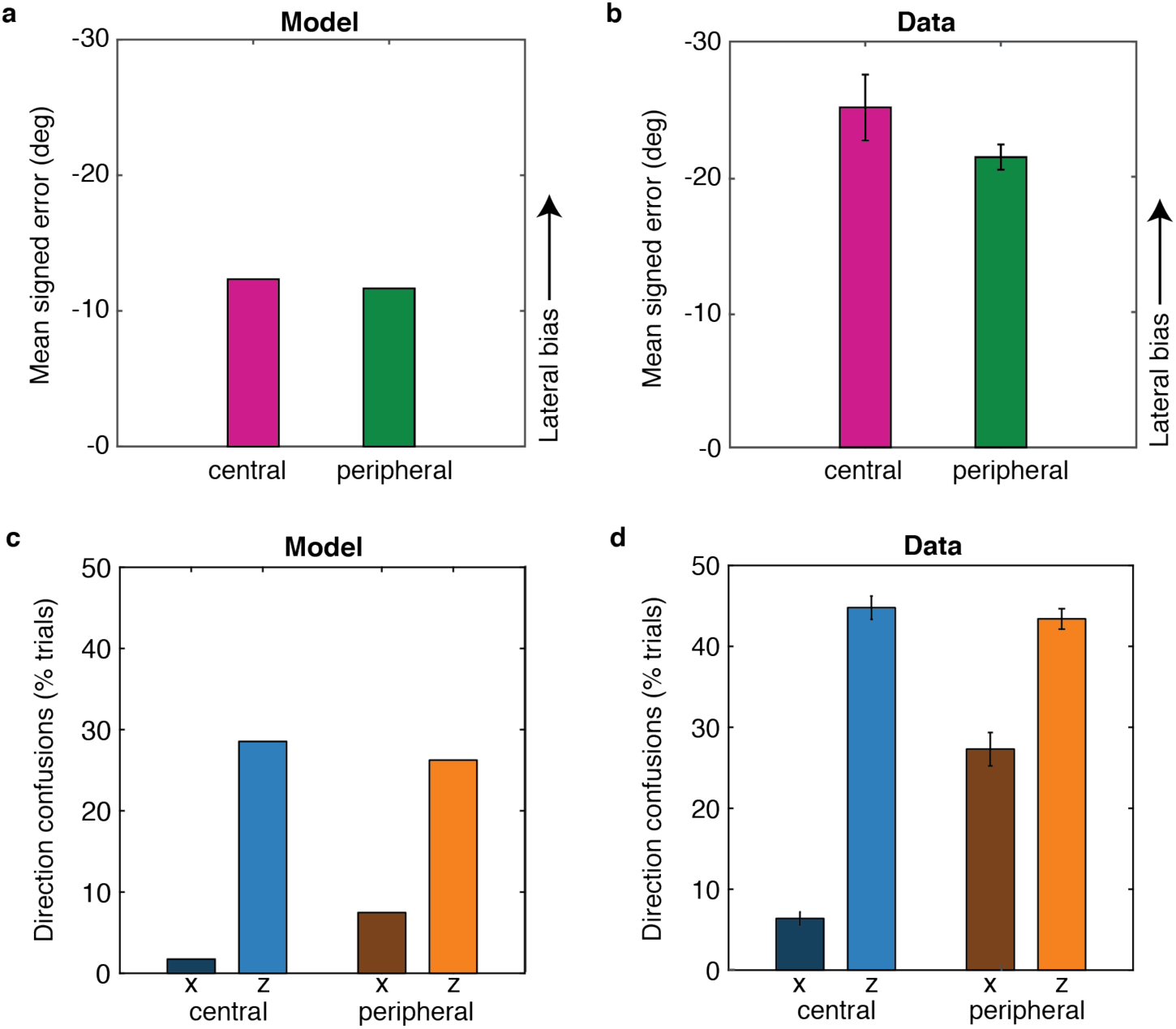
Lateral bias and direction confusions in central and peripheral locations. **(a,b)** Predictions of the model (a) and experimental results for stimuli present in the midsagittal plane (center) and 20 deg to the left or right (periphery). **(c,d)** Predictions of the model (c) and experimental results (d) for lateral motion confusions (x) and motion-in-depth confusions (z). Error bars correspond to +/-1 SEM.

A paired-sample t-test on the data from Experiment 3 revealed no difference in lateral bias in response to targets located at the center and at the periphery (t(20)=-1.45, p=.16). However, on average the lateral bias decreased for peripheral targets by 3.7 deg. There was a small decrease in motion-in-depth direction confusion at the peripheral locations of ~1.38% of trials on average, but this difference was also not significant (t(20)=0.78, p=.44). By contrast, there was a substantial and significant increase in lateral motion direction confusion at the peripheral locations of 20.9% on average (t(20)=-10.82, p<0.001). Thus, consistent with the model predictions, confusions of the lateral motion (x) component of 3D motion stimuli increased significantly with an increase in stimulus eccentricity, whereas confusions for motion-in-depth and the lateral bias did not.

## Discussion

We have presented a Bayesian model of 3D motion perception that predicts systematic errors in perceived motion direction, including a lateral bias, a surprising tendency to report approaching motion as receding and vice versa, and a dependency of these errors on viewing distance, contrast, and eccentricity. We tested these predictions in a VR environment where monocular, binocular, and head motion cues to 3D motion were available, and established that the errors persist under such conditions. Thus, our results demonstrate that uncertainty in retinal velocity signals, coupled with a prior for slow motion and simple geometric considerations, accounts for a number of motion perception phenomena in the three-dimensional world. Our model provides a framework through which to understand errors in 3D motion perception at arbitrary locations, and further supports the idea that visual perception can be accurately and parsimoniously modeled as a process of probabilistic inference.

### Previous Bayesian Models of Motion Perception

This work extends a line of Bayesian models that account for errors in motion perception for stimuli presented in the fronto-parallel plane (Yuille & Grzywacz, 1988; Weiss et al., 2002, Stocker & Simoncelli, 2006). Critically, these models make the assumption that motion percepts are perturbed by the uncertainty in the sensory input combined with a prior for slow motion.

Why would observers employ a prior for slow speeds in the world? A slow motion prior presumably reflects the fact that objects in the world are most likely stationary, and if moving are more likely to move slowly rather than quickly. This prior would thus have to disregard the contributions of eye, head, and body motion to the visual input. Nonetheless, even during head-free fixation, it has been shown that retinal velocity signals are biased towards slower speeds (Aytekin, Victor, & Rucci, 2014). Thus, there is both strong theoretical and experimental evidence that a slow-motion prior would be adaptive for humans.

Two groups have previously extended motion perception models to account for errors in the perception of 3D motion based on binocular cues (Lages, 2006; Welchman et al., 2008). The model proposed by Welchman provides an account for the lateral bias and predicts an effect of viewing distance. However, since the derivation relies on the small angle approximation, this model supports the idea that the perception of motion-in-depth is inherently less reliable than equivalent lateral motion. While this is generally the case for motion in the midsagittal plane, we show that this is by no means a general property of motion-in-depth throughout the visual space. A second added contribution of the current work is to derive the full sampling distribution of the MAP, and therefore account for trial-to-trial variability in judgments of motion direction. Examination of trial-to-trial variability in the experimental data reveals that, in addition to a well-documented lateral bias for motion moving towards the head, observers systematically misreport direction of motion-in-depth such that they judge approaching motion as receding, and vice versa.

While the Welchman model relied on retinal velocity cues, the model proposed by Lages considered the separate contributions of two binocular cues to 3D motion – interocular velocity differences (IOVD) and changing binocular disparity (CD) signals. The Lages study concluded that disparity rather than velocity processing introduced the lateral perceptual bias. However, the Lages model assumed that prior assumptions operate in retinal coordinates (Lages, 2006; Lages & Heron, 2008). Here we have assumed that the combination of the prior and the likelihood takes place in world coordinates. While these assumptions are essentially equivalent for predicting percepts in the fronto-parallel plane, they can produce different predictions for motion-in-depth, depending on whether binocular disparity or binocular motion cues are assumed to be the key visual cue (Lages, 2006). In particular, a model based on combining a prior for motion in retinal coordinates does not predict a lateral bias in 3D motion perception (Lages, 2006).

Interestingly, in the experiments reported in Lages (2006), performance in response to both approaching and receding motion was measured, providing an opportunity to observe the misreports of motion-in-depth reported here. However, in the data analysis, misreports of the depth direction were treated as indications that participants were unable to do the task (if occurring on a large proportion of trials) or ‘bad’ trials in which participants did not see the stimulus. Therefore, such misreports were not treated as a meaningful feature of visual processing until now.

What is the natural coordinate system in which to formulate a Bayesian perceptual model? We would argue that combining the prior and likelihood in world coordinates makes the most sense because it is ultimately motion in the world and not motion on the retina that is relevant to an organism. Thus, the prior and likelihood should be combined in world coordinates to generate a posterior distribution that represents the best-guess for motion in the world, not motion on the retina. However, the extension of a prior for slow speeds into a probability distribution over 3D velocities does not have a single solution. For the current model, we assumed (as has been done previously) that the prior distribution (as well as the likelihood and posterior) is represented in a Cartesian world-space over *x’* and *z’*, where motions towards/away and left/right are continuous with each other (i.e., positive and negative arms of the same axis; see **Figure 7a** and **b** heatmaps). This type of coordinate system is necessary in order for the model to predict the prevalence of direction confusions in depth because the resulting posterior distribution often straddles the *z’*=0 but not the *x’*=0 line. From a purely computation perspective, it would be reasonable to consider that the probabilities of motion trajectories might be represented in terms of polar direction/speed. But in such a coordinate system, it is unclear if the same pattern of direction confusions would result. The clear match between the direction confusion predictions of our model and the experimental data provide strong support that the current model captures essential features that describe the inferences that underlie motion perception.

### Errors in the Real World

The errors predicted by the current model will no doubt be most apparent in the real world under demanding conditions, such as when there is limited time, or poor visibility (Pretto, Bresciani, Rainer, & Bülthoff, 2012; Shrivastava, Hayhoe, Pelz, & Mruczek, 2010; Snowden, Stimpson, & Ruddle, 1998). In situations where sensory uncertainty is very low, the model predicts that these perceptual errors will be negligible. It is difficult to quantify what level of sensory uncertainty a person will be subject to at any particular time during day-to-day life under natural viewing conditions. However, we do know that when stimulus contrast is very high (>100% Michelson contrast), the lateral bias can effectively disappear for practiced observers (Fulvio, Rosen, Rokers, 2015). While the motion-in-depth confusions persist longer in the laboratory, we expect that these may be similarly reduced by the presence of additional and more reliable visual cues. In fact, the presence of these systematic errors may provide a way to compare and quantify the performance of different virtual reality display systems, especially those that incorporate less well-understood cues such as predictive head motion, or defocus blur.

### Implications for Neural Processing of Motion

While the current model is perceptual and not mechanistic, our predictions and results are relevant to investigating the neural mechanisms that underlie motion perception. The central role of area MT in the processing of binocular 3D motion signals is now well-established, based on both neuroimaging (Rokers, Cormack, Huk, 2009) and electrophysiology studies (Czuba, Huk, Cormack, & Kohn, 2014; Sanada & DeAngelis, 2014). Our model highlights the fact that both position and binocular speed tuning are essential for inferring the trajectory of a stimulus moving in 3D. Take the case of an object moving directly towards the mid-point between the two eyes. If this object is located in the midsagittal plane, it will cast equal and opposite horizontal velocities in the two eyes. However, if this object has an eccentric location to the left of the midsagittal plane, the velocities cast on the two eyes will not be equal and opposite – they will have opposite signs but the velocity in the left eye will be greater. Thus, the interpretation of an MT neuron’s tuning profile and preference for 3D motion must somehow take location of the stimulus relative to the observer into account, independent of retinotopic location.

When it comes to the slow motion prior, there remains significant debate on how prior assumptions for visual motion factor into the neural computations, and where perceptual biases arise along the visual motion processing pathway. Results from neuroimaging show that responses to 2D motion stimuli can depend on perceived rather than presented speed as early as V1, suggesting that motion priors interact with sensory evidence at the earliest stage of cortical processing (Vintch & Gardner, 2014). However, evidence from electrophysiology has been decidedly more mixed (Pack, Hunter, Born, 2005; Krekelberg, van Wezel, & Albright, 2006; Livingstone & Conway, 2007). Since the biases for the lateral motion and motion-in-depth components for 3D stimuli have different magnitudes, these differences provide an additional signature for determining whether the responses of particular neuronal populations are driven by the stimulus or the percept.

### Conclusion

Understanding how Bayesian inference plays out during natural vision and natural behavior requires not only characterizing the prior assumptions of an observer, but also having a deep understanding of the sensory signals available at a given point in time. The current model predicts perceived 3D motion under a wide range of scenarios and viewing conditions, and in doing so provides a parsimonious account of multiple seemingly disparate perceptual errors.

## Acknowledgements

The authors would like to thank Padmadevan Chettiar for technical support, Michelle Wang and Darwin Romulus for assistance with data collection, and Joe Austerweil for comments on a previous version of this manuscript. E.A.C was supported by Oculus, Microsoft, and Samsung.

